# Rapid analysis of strigolactone receptor activity in a *Nicotiana benthamiana dwarf14* mutant

**DOI:** 10.1101/2021.05.11.443507

**Authors:** Alexandra R.F. White, Jose A. Mendez, Aashima Khosla, David C. Nelson

## Abstract

DWARF14 (D14) is an ɑ/β-hydrolase and receptor for the plant hormone strigolactone (SL) in angiosperms. Upon SL perception, D14 works with MORE AXILLARY GROWTH2 (MAX2) to trigger polyubiquitination and degradation of DWARF53(D53)-type proteins in the SUPPRESSOR OF MAX2 1-LIKE (SMXL) family. We used CRISPR-Cas9 to generate knockout alleles of the two homoeologous *D14* genes in the *Nicotiana benthamiana* genome. The *Nbd14a,b* double mutant had several phenotypes that are consistent with the loss of SL perception in other plants, including increased axillary bud outgrowth, reduced height, shortened petioles, and smaller leaves. A ratiometric fluorescent reporter system was used to monitor degradation of SMXL7 from *Arabidopsis thaliana* (AtSMXL7) after transient expression in *N. benthamiana* and treatment with the strigolactone analog GR24. AtSMXL7 was degraded after treatment with GR24^5DS^, which has the stereochemical configuration of SLs, as well as its enantiomer GR24^*ent*-5DS^ . In *Nbd14a,b* leaves, AtSMXL7 abundance was unaffected by GR24. Transient coexpression of AtD14 with the AtSMXL7 reporter in *Nbd14a,b* restored the degradation response to GR24, but required an active catalytic triad. With this platform, we evaluated the ability of several AtD14 mutants that had not been characterized in plants to target AtSMXL7 for degradation.

## INTRODUCTION

Strigolactones (SLs) are a family of plant hormones derived from β-carotene that have diverse functions in plants (Waters et al., 2017; Machin et al., 2020; Bouwmeester et al., 2021). SLs regulate axillary bud outgrowth (tillering), stem elongation, auxin transport, root elongation, leaf shape, leaf angle, leaf senescence, cambial growth, susceptibility to pathogenic microbes and root-knot nematodes, stomatal closure responses, and drought tolerance (Gomez-Roldan et al., 2008; Umehara et al., 2008; Agusti et al., 2011; Kapulnik et al., 2011; Ruyter-Spira et al., 2011; Scaffidi et al., 2013; Shinohara et al., 2013; Bu et al., 2014; Van Ha et al., 2014; Yamada et al., 2014; Lauressergues et al., 2015; Soundappan et al., 2015; Ueda and Kusaba, 2015; Marzec et al., 2016; Li et al., 2017; Lahari et al., 2019; Nasir et al., 2019; Kalliola et al., 2020; Li et al., 2020; Shindo et al., 2020). SLs are also exuded by roots into the rhizosphere, especially under nutrient-poor conditions. There, SLs stimulate hyphal branching and metabolic activity of arbuscular mycorrhizal (AM) fungi, promoting beneficial symbiotic interactions with the host plant (Akiyama et al., 2005; Besserer et al., 2006; Besserer et al., 2008; Kretzschmar et al., 2012; Kobae et al., 2018).

SL perception in flowering plants is mediated by the ɑ/β-hydrolase DWARF14 (D14)/DECREASED APICAL DOMINANCE (DAD2)/RAMOSUS3 (RMS3) (Arite et al., 2009; Hamiaux et al., 2012; Waters et al., 2012; de Saint Germain et al., 2016). Upon activation by SL, D14 associates with the F-box protein MORE AXILLARY GROWTH2 (MAX2)/DWARF3(D3), which acts as an adapter component of an SCF (Skp1–Cullin–F-box) E3 ubiquitin ligase complex. Activated D14 also associates with a subset of proteins in the SUPPRESSOR OF MAX2 1-LIKE (SMXL) family that are known as DWARF53 (D53) in rice and petunia, and SMXL6, SMXL7, and SMXL8 in *Arabidopsis thaliana*. This leads to the polyubiquitination of D53-type SMXLs by SCF^MAX2^ and their degradation by the 26S proteasome (Jiang et al., 2013; Zhou et al., 2013; Soundappan et al., 2015; Wang et al., 2015a; Yao et al., 2016; Shabek et al., 2018; Lee et al., 2020). In *Arabidopsis thaliana*, D14 is also able to associate with SMXL2 and target it for degradation when a SL analog is applied (Wang et al., 2020).

A very similar signaling mechanism is used by karrikins (KARs), a class of plant growth regulators found in smoke. KAR signaling requires SCF^MAX2^ and an ancient paralog of D14 known as KARRIKIN INSENSITIVE2 (KAI2)/HYPOSENSITIVE TO LIGHT (HTL) (Nelson et al., 2011; Sun and Ni, 2011; Waters et al., 2012). KAI2-SCF^MAX2^ targets SMAX1 and its close paralog SMXL2 for degradation (Stanga et al., 2013; Stanga et al., 2016; Khosla et al., 2020a; Wang et al., 2020; Zheng et al., 2020). This pathway regulates seed germination, hypocotyl/mesocotyl elongation, seedling responses to light, leaf shape, cuticle development, drought tolerance, root skewing, root hair density and elongation, and the capacity for arbuscular mycorrhizal symbiosis (Shen et al., 2007; Nelson et al., 2009; Nelson et al., 2010; Sun and Ni, 2011; Stanga et al., 2013; Gutjahr et al., 2015; Soundappan et al., 2015; Stanga et al., 2016; Li et al., 2017; Swarbreck et al., 2019; Villaécija-Aguilar et al., 2019; Bunsick et al., 2020; Carbonnel et al., 2020; Choi et al., 2020; Li et al., 2020; Zheng et al., 2020).

*D14* has highly conserved roles in angiosperms. This has been demonstrated through analysis of *d14* mutants in petunia (*Petunia hybrida*), rice (*Orzya sativa*), *Arabidopsis thaliana*, canola (*Brassica napus*), pea (*Pisum sativum*), barley (*Hordeum vulgare*), hexaploid wheat (*Triticum aestivum*), *Medicago truncatula*, and *Lotus japonicus*, as well as RNAi knockdown of *D14* in soybean hairy roots (Arite et al., 2009; Hamiaux et al., 2012; Waters et al., 2012; Lauressergues et al., 2015; Marzec et al., 2016; de Saint Germain et al., 2016; Carbonnel et al., 2019; Ahmad et al., 2020; Liu et al., 2021; Stanic et al., 2021). *D14* orthologs from cotton, *Populus trichocarpa*, and chrysanthemum have also been studied indirectly through cross-species complementation of an *Arabidopsis d14* (*Atd14*) mutant (Wen et al., 2015; Zheng et al., 2016; Wang et al., 2019). This approach enables *in vivo* analysis of gene function for species that have fewer genetic resources available or are less tractable to genetic studies than the major model plant systems (e.g. species lacking insertion/TILLING mutant collections, effective transformation methods, etc.). For example, this method has been used to identify strigolactone receptors in root parasitic plants that arose from neofunctionalization of *KAI2*/*HTL* paralogs (Conn et al., 2015; Toh et al., 2015; de Saint Germain et al., 2020).

The utility of the cross-species complementation approach is limited by the compatibility of the transgene product of interest with its non-cognate cellular environment. For example, some *MAX2* and *KAI2/HTL* transgenes from petunia; the bryophytes *Selaginella moellendorffii, Marchantia polymorpha*, and *Physcomitrium* (formerly *Physcomitrella*) *patens*; and the parasitic plant *Striga hermonthica* are nonfunctional or only partially functional in *Arabidopsis* (Drummond et al., 2011; Liu et al., 2014; Conn et al., 2015; Toh et al., 2015). This does not necessarily mean that these genes have reduced function in their native context; instead, the proteins they encode may not be able to interact well with *Arabidopsis* orthologs of their signaling partners (Khosla and Nelson, 2016). For example, at least two *Striga hermonthica* KAI2/HTL proteins that are inactive in *Arabidopsis* can bind SL sensitively *in vitro* but cannot interact with *Arabidopsis* MAX2 (AtMAX2), or for that matter ShMAX2 (Wang et al., 2021). Conversely, *Striga hermonthica* HTL7 (ShHTL7) causes *Arabidopsis* seed to germinate in the presence of picomolar SL. This response is several orders of magnitude lower than that conferred by ShHTL proteins with similar SL affinities *in vitro*, and is likely due to the unusually high affinity of ShHTL7 for AtMAX2 (Toh et al., 2015; Tsuchiya et al., 2015; Uraguchi et al., 2018; Wang et al., 2021). Another disadvantage of the cross-species complementation approach to investigate gene function is that it typically takes several generations to obtain homozygous transgenic lines that are suitable to study. Therefore, methods to evaluate plant gene function *ex situ* that are fast and also allow the co-introduction of compatible transgene partners are desirable.

We previously developed a ratiometric reporter system (pRATIO) that can monitor changes in the relative abundance of a transiently expressed target protein compared to a reference protein (Khosla et al., 2020b). In this Gateway-compatible system, a gene of interest (target) is translationally fused to a fluorescent or bioluminescent reporter. The target is co-transcribed with a reference gene that also encodes a fluorescent or bioluminescent reporter (Khosla et al., 2020b). A modified “self-cleaving” 2A peptide derived from foot-and-mouth disease virus is encoded between the two genes, causing the target and reference proteins to be translated separately. This approach enables multicistronic, stoichiometric expression in eukaryotes (Luke et al., 2010).

The pRATIO system has been applied successfully in *Nicotiana benthamiana*, a native Australian species that is closely related to tobacco (*N. tabacum*), to investigate degradation of *Arabidopsis* SMAX1, SMXL7, and KAI2 (Bally et al., 2018; Khosla et al., 2020a; Khosla et al., 2020b). When transiently expressed in *N. benthamiana* leaves, the *Arabidopsis* SMXL7 (AtSMXL7) ratiometric reporter is degraded after treatment with *rac*-GR24, a racemic mixture of the synthetic SL analog GR24^5DS^ and its enantiomer, GR24^ent-5DS^ . In contrast, KAR_1_ or KAR_2_ treatments, which are expected to activate KAI2, do not affect AtSMXL7 stability (Khosla et al., 2020a). This is consistent with prior studies of rice D53 and *Arabidopsis* SMXL6, SMXL7, and SMXL8 degradation in response to GR24, but not KAR_1_ treatments (Jiang et al., 2013; Zhou et al., 2013; Wang et al., 2015a). It also indicates that a GR24-responsive receptor(s) in *N. benthamiana* is able to target AtSMXL7 for degradation.

GR24-induced degradation of D53-type SMXLs is dependent on *D14* in rice and *Arabidopsis* (Jiang et al., 2013; Zhou et al., 2013; Wang et al., 2015a; Samodelov et al., 2016). Genetic and evolutionary evidence also support the idea that AtSMXL7 is targeted by AtD14 and not AtKAI2 (Soundappan et al., 2015; Waters et al., 2015; Machin et al., 2020). Therefore, the AtSMXL7 degradation response is likely mediated by one or both of the two nearly identical *D14* homoeologs found in the *N. benthamiana* genome. However, the possibility has been raised that there can be crosstalk between KAI2 and SMXL6, SMXL7, and SMXL8 in the regulation of *Arabidopsis* root skewing (Swarbreck et al., 2019). The *N. benthamiana* genome contains four *KAI2* paralogs that are likely to encode functional proteins. Also, KAI2 proteins are not limited to KAR perception. *Arabidopsis* KAI2 is activated by GR24^*ent*-5DS^, while in parasitic plants some evolutionarily “divergent” KAI2 (KAI2d) proteins are able to perceive GR24^5DS^ and natural SLs (Conn et al., 2015; Toh et al., 2015; Tsuchiya et al., 2015; de Saint Germain et al., 2020; Nelson, 2021). Therefore, it is not clear whether any NbKAI2 proteins might contribute to *rac*-GR24-induced degradation of AtSMXL7 in *N. benthamiana*.

We reasoned that transient co-expression in *N. benthamiana* could provide a way to rapidly evaluate the ability of D14 variants to target a SMXL7 ratiometric reporter for degradation. This could enable medium-throughput screens for mutations that affect D14 signaling activity or its protein-protein interactions with SMXL7. Pairs of D14-SMXL7 proteins from other species could also be evaluated as long as compatibility with *N. benthamiana* MAX2 is maintained. For this approach to be most effective, however, the endogenous SL receptors in *N. benthamiana* would need to be removed. Therefore, we used CRISPR-Cas9 to knock out *N. benthamiana D14a* and *D14b*. We evaluate the combined roles of these genes in *N. benthamiana* shoot development. We also demonstrate that this mutant background can be used to analyze the capacity of *Arabidopsis* D14 variants to degrade AtSMXL7.

## RESULTS

### Knockout of two *D14* genes in *Nicotiana benthamiana* with CRISPR-Cas9

*Nicotiana benthamiana* is an allotetraploid that carries two *D14* homoeologs in its genome (Figure 1A). The coding sequences of *NbD14a* and *NbD14b* are 97% identical at the nucleotide level, resulting in only three amino acid differences (Ala/Thr84, Leu/Ile119, Ala/Thr257) between the 267-aa proteins. We selected a pair of Cas9-compatible gRNAs that would simultaneously target both *NbD14* genes in each of their two exons (Figure 1). These gRNAs were cloned into an egg cell-specific promoter-controlled CRISPR-Cas9 vector that was originally developed for use in *Arabidopsis thaliana* (Wang et al., 2015b). This construct was stably introduced into wild-type *N. benthamiana* through *Agrobacterium tumefaciens*-mediated transformation. We identified a homozygous *Nbd14a Nbd14b* double mutant (hereafter, *Nbd14a,b*) in the T_0_ generation and subsequently isolated a line free of the CRISPR-Cas9 transgene. The *Nbd14a-1* allele is composed of two mutations: the first is a 14 bp deletion in exon 1 that results in a frameshift after Arg109 and premature truncation of the protein, and the second is a 1 bp deletion in exon 2 (Figure 1B). The *Nbd14b-1* allele is a 1 bp deletion at the same position in exon 2, which causes a frameshift after Cys209 (Figure 1B). This results in loss of the Asp217 and His246 residues of the catalytic triad in addition to many other amino acids. Both Asp217 and His246 are required for SL hydrolysis by *Arabidopsis* D14 *in vitro*, and the catalytic histidine residue is necessary for D14 activity *in planta* (Seto et al., 2019). Therefore, both *d14* alleles are expected to cause a complete loss-of-function.

**Figure 1.**
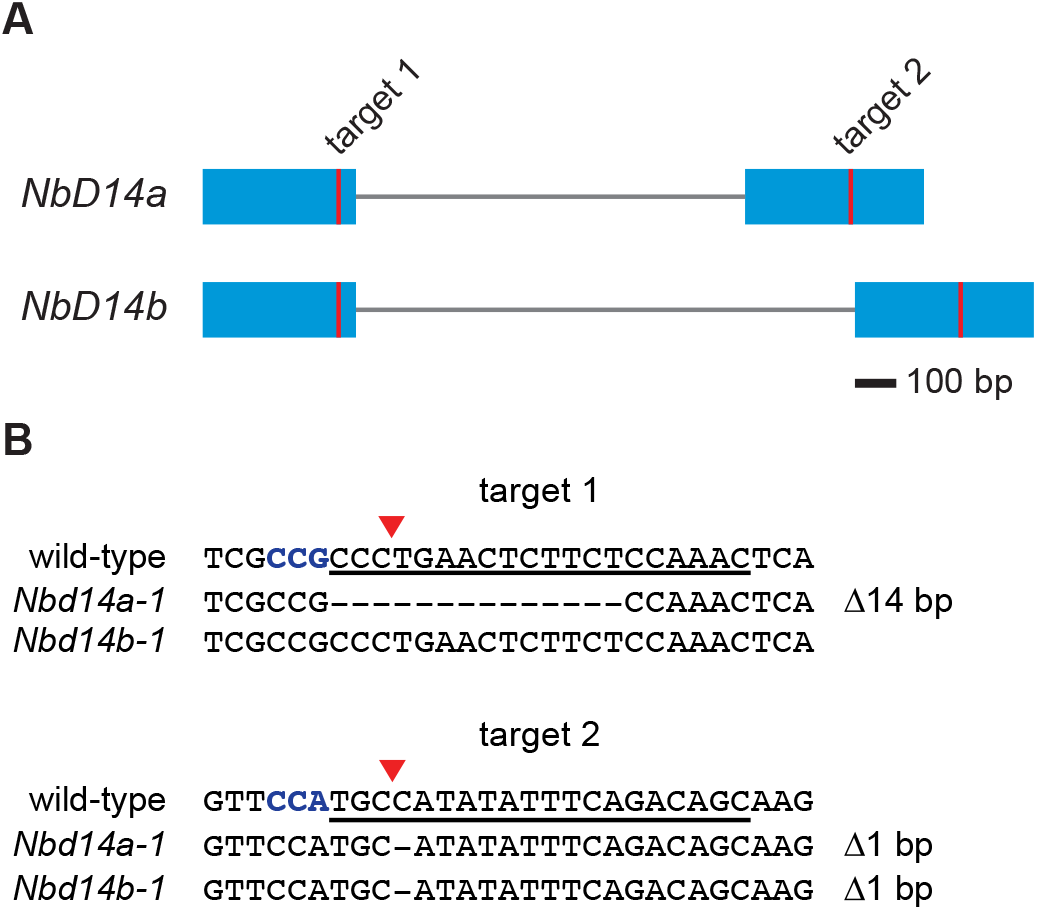
Mutation of two *D14* genes in *Nicotiana benthamiana* with CRISPR-Cas9. (A) Diagram of *D14a* and *D14b* genes in *N. benthamiana*. Blue boxes represent exons. Vertical red lines indicate gRNA target sites. (B) Sequences of Cas9-induced frameshift alleles of *NbD14a* and *NbD14b*. gRNA target sequence is underlined. Protospacer adjacent motif (PAM) sequence (5’-NGG-3’) is indicated in bold blue font. Red triangles denote predicted Cas9 cleavage sites.

### *N. benthamiana d14a,b* has increased shoot branching and altered leaf shape

Loss-of-function mutations in *D14* cause increased branching/tillering and semi-dwarf stature in rice, barley, *Arabidopsis*, canola, petunia, pea, *Medicago truncatula*, and *Lotus japonicus* (Arumingtyas et al., 1992; Napoli and Ruehle, 1996; Arite et al., 2009; Hamiaux et al., 2012; Waters et al., 2012; Lauressergues et al., 2015; Marzec et al., 2016; de Saint Germain et al., 2016; Carbonnel et al., 2019; Stanic et al., 2021). To determine whether *D14* has a similar role in *N. benthamiana*, we examined the shoot architecture of 5-week old plants grown under greenhouse conditions. To our complete lack of surprise, *Nbd14a,b* plants had a more compact, “bushy” shoot architecture compared to wild-type (Figure 2A). We measured outgrowth of the first nine axillary buds of each plant and found that the six most basal axillary shoots were significantly longer in *Nbd14a,b* than wild-type (Figure 2A, B). At the younger, apical nodes, bud outgrowth was reduced and there was no significant difference in the lengths of *Nbd14a,b* and wild-type axillary shoots. This pattern of basitonic development, in which basal branches show more vigorous outgrowth than the apical branches, is also found in the *decreased apical dominance* (*dad*) mutants of *Petunia hybrida*, a related solanaceous species (Napoli and Ruehle, 1996). Consistent with the semi-dwarf phenotype of *d14* mutants in other angiosperms, we also observed that the height of *d14a,b* plants was reduced significantly compared to wild-type (Figure 2A, C).

**Figure 2.**
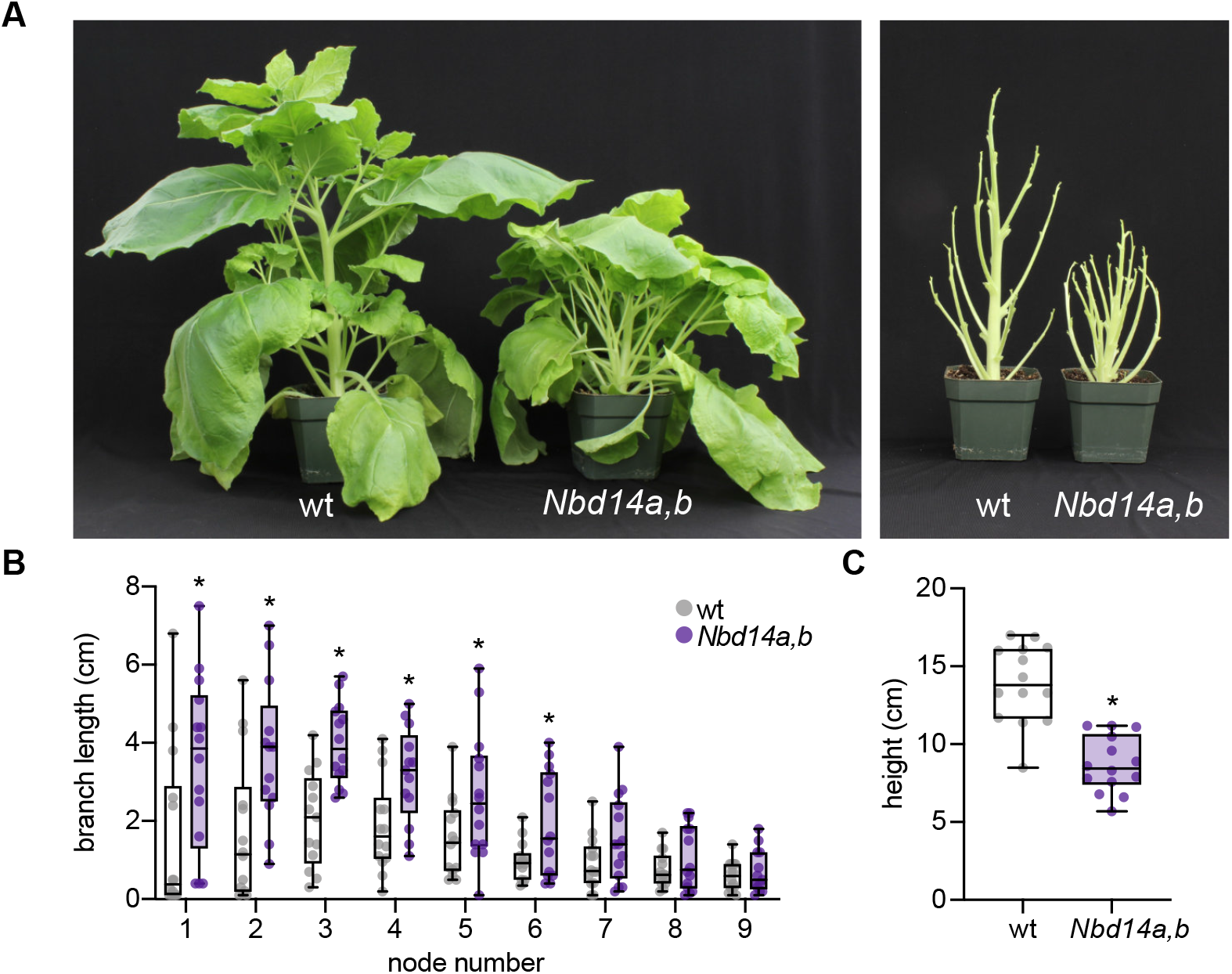
*Nbd14a,b* has more axillary bud outgrowth and reduced height. (A) Photographs of 10-week old wt and *Nbd14a,b N. benthamiana* plants with (right) and without (left) leaves. (B) Bud outgrowth of the first nine axillary nodes of 58-day old wt and *Nbd14a,b* plants. (C) Primary shoot height of wt and 58-day old *Nbd14a,b*. Box plots show median with 25th and 75th percentiles, and whiskers represent minimum and maximum values; n=10-12 plants; *, p <0.05, unpaired t-test with Welch correction, comparison of *Nbd14a,b* to wt.

The petioles of *Arabidopsis d14* leaves have substantially reduced length compared to wild-type. In addition, the length and width of *d14* leaves are reduced, resulting in a smaller and more rounded blade shape overall (Waters et al., 2012; Scaffidi et al., 2013; Soundappan et al., 2015). In *Medicago truncatula, d14* leaflets have increased “solidity” due to increased but shallower serrations at the leaflet margin (Lauressergues et al., 2015). These observations led us to examine leaf morphology in *Nbd14a,b*. We measured the petiole length, blade length, and blade width of the subtending leaf for each of the first nine axillary buds (Figure 3). The petioles of *Nbd14a,b* leaves one (first-emerged, basal) through six were significantly shorter than wild-type (Figure 3A). Blade length and width were also reduced significantly for most of the six oldest *Nbd14a,b* leaves (Figure 3B, C). The most apical leaves had similar dimensions in *d14a,b* and wild-type, at least at this time point. Together, these data indicate that loss of *NbD14a* and *NbD14b* alters the shoot architecture and leaf morphology of *N. benthamiana* similarly to *d14* mutants in other angiosperms.

**Figure 3.**
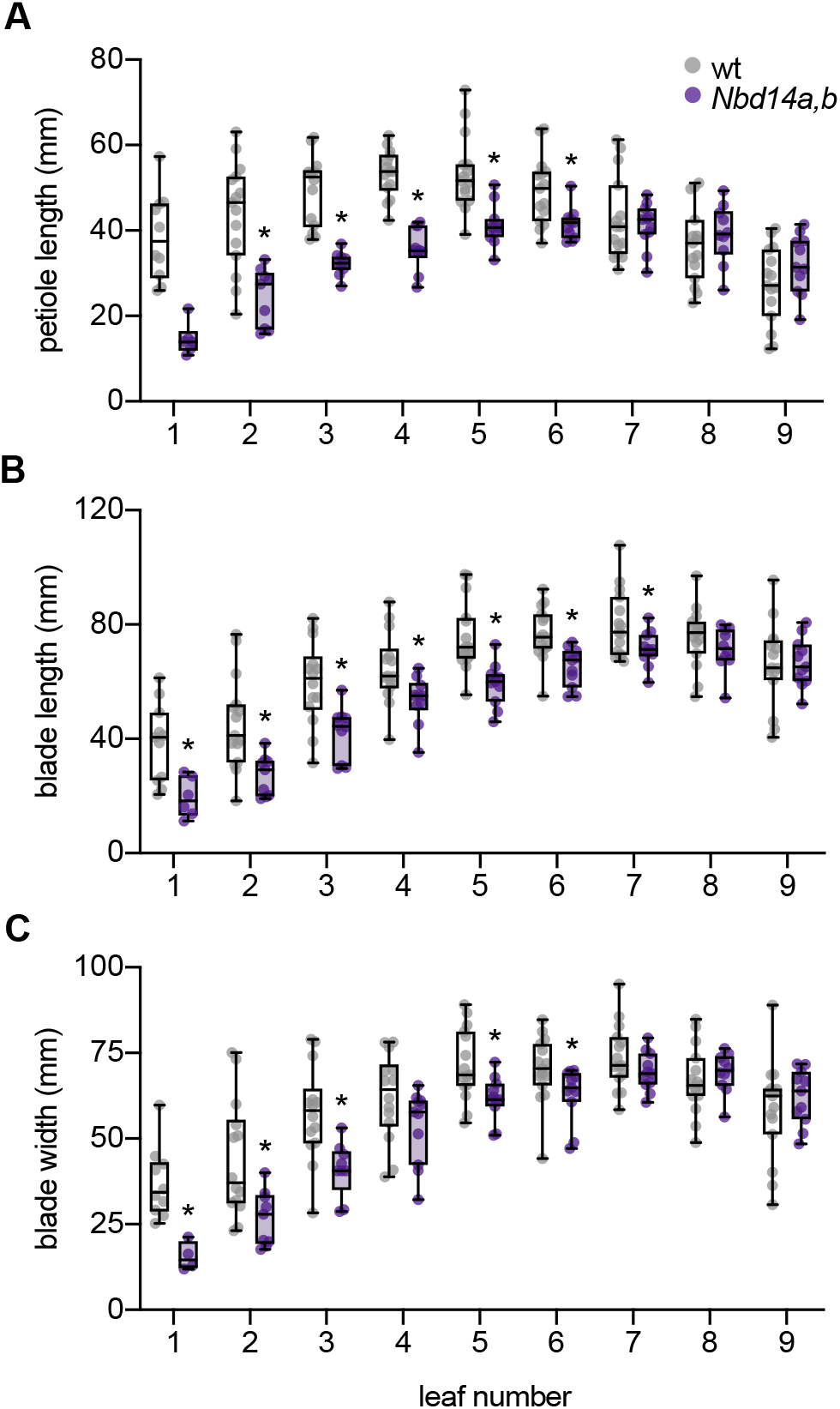
*Nbd14a,b* has smaller leaves and petioles. (A) Petiole lengths (B) blade lengths, and (C) blade widths of the first nine leaves of 58-day old wt and *Nbd14a,b* plants. Box plots show median with 25th and 75th percentiles, and whiskers represent minimum and maximum values; n=10-12 plants; *, p <0.05, unpaired t-test with Welch correction, comparison of *Nbd14a,b* to wt.

### GR24-stimulated degradation of AtSMXL7 in *N. benthamiana* requires *NbD14a,b*

We previously used a ratiometric reporter system (pRATIO3212) to show that AtSMXL7 expressed in wild-type *N. benthamiana* leaves is degraded after treatment with *rac*-GR24, but not KAR_1_ or KAR_2_ (Khosla et al., 2020a). To determine whether it is the GR24^5DS^ or GR24^*ent-*5DS^ component of *rac*-GR24 that triggers AtSMXL7 degradation, we tested the effects of optically pure compounds. Both GR24^5DS^ and GR24^*ent-*5DS^ caused a statistically significant reduction in the ratio of fluorescence signals from the AtSMXL7-mScarlet-I target relative to the reference protein, Venus (Figure 4). However, GR24^5DS^ had a stronger effect on the decline of the SMXL7 reporter than GR24^*ent-*5DS^, at least within the 16 h treatment period.

**Figure 4.**
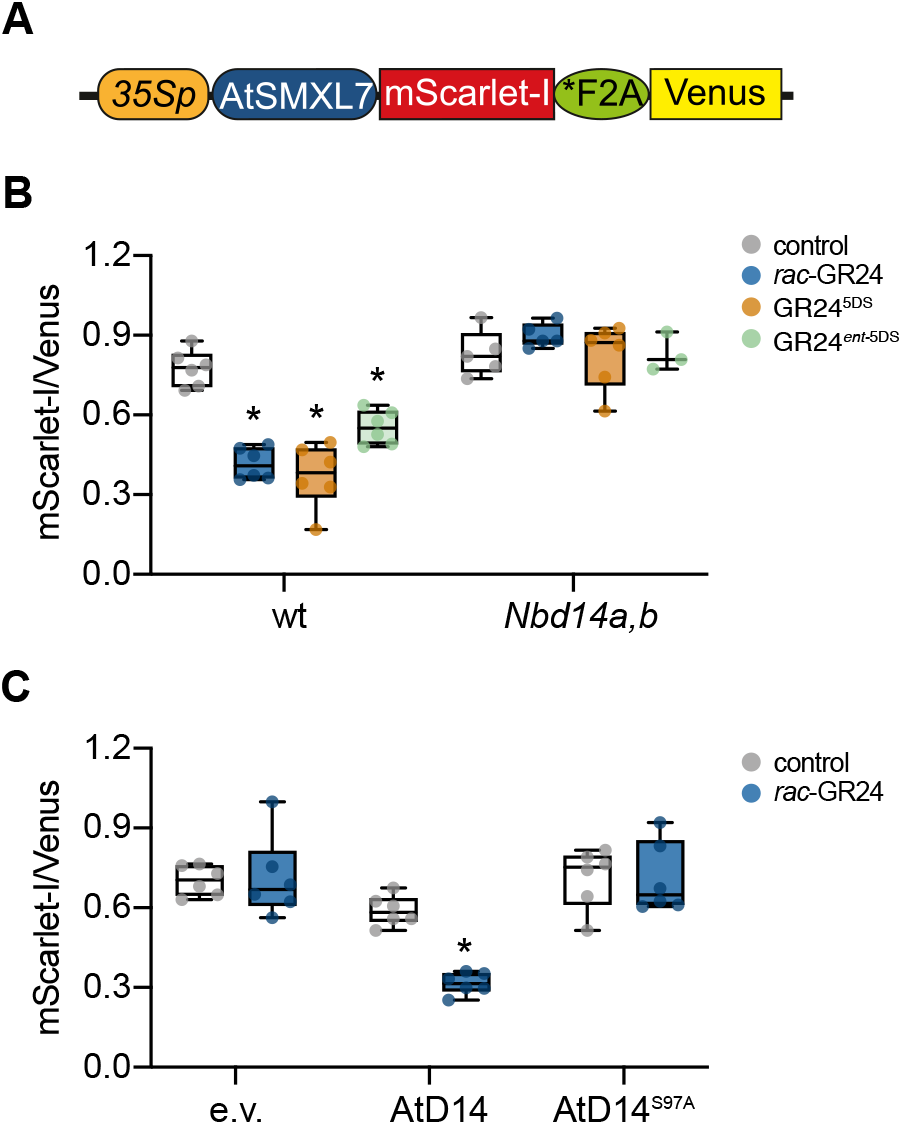
AtSMXL7 degradation in *N. benthamiana* is *NbD14*-dependent. Diagram of the ratiometric AtSMXL7 reporter expressed in pRATIO3212. mScarlet-I is a fluorescent reporter protein translationally fused to the C-terminus of AtSMXL7, *F2A is a modified “self-cleaving” peptide, and Venus is a yellow fluorescent protein used for normalization. Diagram adapted from (Khosla et al., 2020b). (B) Background-corrected AtSMXL7-mScarlet-I to Venus fluorescence in *N. benthamiana* wt and *Nbd14a,b* leaf discs after 16 h treatments with solvent control (0.02% acetone (v/v)), or 10μM *rac*-GR24, GR24^5DS^, or GR24^*ent-*5DS^ . (C) Background-corrected AtSMXL7-mScarlet-I to Venus fluorescence in *Nbd14a,b* leaf discs after 16 h treatments with solvent control (0.02% acetone (v/v)), or 10μM *rac*-GR24. pRATIO3212-AtSMXL7 was coexpressed with pGWB415 empty vector (e.v.), AtD14, or AtD14^S97A^ . Box plots show median with 25th and 75th percentiles, and whiskers represent minimum and maximum values; n=3-6 leaf discs; *, p <0.05, unpaired t-test with Welch correction, comparison of treatment to control.

We investigated whether GR24-induced degradation of AtSMXL7 in *N. benthamiana* is D14-dependent, or if other proteins such as KAI2 might also contribute. AtSMXL7-mScarlet-I abundance was not affected by treatment with *rac*-GR24 or either of the purified GR24 stereoisomers in *Nbd14a,b* leaves (Figure 4B). This strongly suggests that AtSMXL7 degradation after GR24 treatment in *N. benthamiana* is only caused by NbD14 proteins. Considering the prior experiment (Figure 4A), this result further implies that NbD14 proteins are more responsive to GR24^5DS^, which shares a stereochemical configuration with naturally occurring SLs, than to GR24^*ent*-5DS^ . *Arabidopsis* D14 shows a similar preference for GR24^5DS^ (Scaffidi et al., 2014; Flematti et al., 2016; Samodelov et al., 2016).

We next tested whether co-transformation of AtD14 could restore GR24-induced degradation of the AtSMXL7 reporter to the *Nbd14a,b* mutant. As negative controls, we compared the effects of an empty vector (pGWB415) and a AtD14^S97A^ mutant on AtSMXL7 degradation. Like many other ɑ/β-hydrolases, D14 has a highly conserved Ser-His-Asp catalytic triad that is necessary for its enzymatic activity. AtD14^S97A^ does not hydrolyze SL and is nonfunctional in plants (Hamiaux et al., 2012; Abe et al., 2014; Seto et al., 2019). Treatment with 10 µM *rac*-GR24 caused a decrease (p<0.01, Welch’s t-test) in the AtSMXL7 target-to-reference ratio when *35S:AtD14* was co-transformed, but had no effect on samples co-transformed with empty vector or *35S:AtD14*^*S97A*^ (Figure 4C). This demonstrated that transient expression of *AtD14* could rescue SL signaling in *Nbd14a,b*. It also raised the possibility that this approach could be used to evaluate the ability of different D14 variants to trigger SMXL7 degradation.

### A rapid assay for the induction of AtSMXL7 degradation by AtD14 mutants

A recent study identified several amino acid substitutions that affect yeast two-hybrid interactions of *Petunia x hybrida* DAD2 (PhDAD2, a D14 ortholog) with PhMAX2A and/or the SMXL7 ortholog PhD53A (Lee et al., 2020). An F135A substitution enhanced PhDAD2 interactions with PhD53A in the absence of *rac*-GR24, but did not affect interactions with PhMAX2A. By contrast, N242I enhanced PhDAD2 interactions with PhMAX2A, but not PhD53A, in the absence of *rac*-GR24. When these substitutions were combined, the PhDAD2 mutant protein showed enhanced interactions with both PhD53A and PhMAX2A that were not further stimulated by *rac*-GR24. A third substitution, D166A, disrupted PhDAD2 interactions with PhMAX2A but not PhD53A (Lee et al., 2020). Based on these results, PhDAD2^F135A^ and PhDAD2^N242I^ might be expected to have hypersensitive responses to SL, whereas PhDAD2^D166A^ may be insensitive to SL. Because these PhDAD2 variants were not tested in plants, however, it is possible that some of the altered interactions are specific to yeast two-hybrid and do not translate to effects on SL signaling activity.

We reasoned that the *Nbd14a,b* mutant could provide a fast assay of D14 signaling activity that complements approaches such as yeast two-hybrid. We synthesized amino acid substitutions in AtD14 that were equivalent to those previously characterized in PhDAD2: F136A, K243I, F136A/K243I, and D167A. We then tested the ability of these AtD14 variants to degrade an AtSMXL7 ratiometric reporter after 25 nM and 10 µM *rac*-GR24 treatments when co-expressed in *Nbd14a,b* leaves. AtD14^F136A^ showed a stronger response to 25 nM rac-GR24 than wild-type AtD14 (Figure 5A). AtD14^K243I^ caused lower abundance of the AtSMXL7 reporter in the absence of rac-GR24 treatment than the other D14 proteins, suggesting it may have semi-constitutive SL signaling activity or is more sensitive to endogenous SLs in *N. benthamiana*. The AtD14^F136A/K243I^ double mutant produced unexpected results. In contrast to the yeast two-hybrid results, this mutant protein showed a positive SL signaling response to *rac*-GR24 (Figure 5A). AtD14^D167A^ also showed a positive, albeit reduced, response to *rac*-GR24 that would not necessarily have been predicted from the yeast two-hybrid assays (Figure 5B).

**Figure 5.**
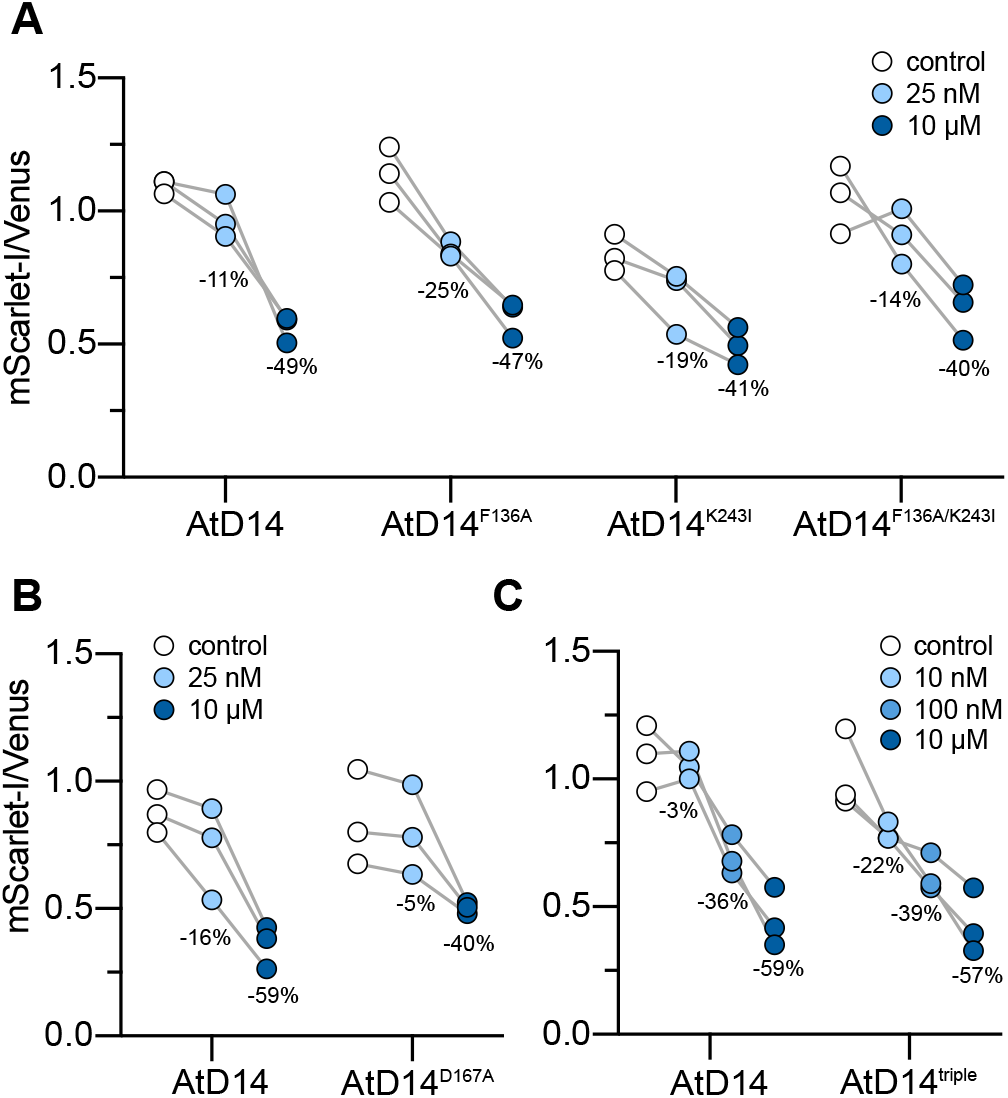
AtSMXL7 degradation is altered when co-expressed with AtD14 variants. Background-corrected AtSMXL7-mScarlet-I to Venus fluorescence in *Nbd14a,b* leaf discs after 16 h treatments with solvent control (0.02% acetone (v/v)), 10 nM, 25 nM, 100 nM, or 10 μM *rac*-GR24. pRATIO3212-AtSMXL7 was coexpressed with pGWB415-AtD14, (A) pGWB415-AtD14^F136A^, -AtD14^K243I^, or -AtD14^F136A/K243I^ ; (B) pGWB415-AtD14^D167A^ ; or (C) AtD14^F159T/A163L/G165A^ (AtD14^triple^). Each point represents the mean of n=3-6 treated discs from an individually transformed leaf. Paired data are linked by lines. Percentages indicate mean percent reduction in mScarlet-I/Venus ratio of each treatment compared to control.

Another recent study examined the basis of the exceptionally sensitive SL response of *Striga hermonthica* HTL7 (ShHTL7), a member of an evolutionarily divergent class of KAI2 proteins (KAI2d) found in root parasitic plants (Wang et al., 2021). When expressed in *Arabidopsis thaliana*, ShHTL7 confers germination responses to picomolar concentrations of SL (Toh et al., 2015). Paralogous KAI2d proteins, such as ShHTL6, show similar affinities to SL *in vitro* as ShHTL7 but do not produce such sensitive SL responses in Arabidopsis (Toh et al., 2015; Wang et al., 2021). At least one likely reason that ShHTL7 is so effective in Arabidopsis is that it has an unusually high affinity for MAX2 upon activation. This high affinity for MAX2 was recreated in ShHTL6 through five amino acid substitutions to mimic ShHTL7 identities: F157T, M161L, G163A, S180N, and I181M (Wang et al., 2021). It must be noted, however, that the effect of these substitutions was only tested through *in vitro* pull-downs, and not in plants.

We explored whether creating similar substitutions in AtD14 might enhance its response to SL *in vivo*. We synthesized an AtD14 variant with three substitutions (F159T, A163L, and G165A), as the equivalent residues of Asn180 and Met181 in AtD14 already shared the same identities as ShHTL7. When co-transformed into *Nbd14a,b* leaves, AtD14^F159T/A163L/G165A^ (AtD14^triple^) caused more degradation of AtSMXL7 in the presence of 10 nM *rac*-GR24 than wild-type AtD14 (Figure 5B). This suggested that the F159T/A163L/G165A substitutions may enhance SL signaling activity by D14. Further investigation of stable transgenic plant lines in Arabidopsis and biochemical assays for interactions with MAX2 will be required to determine whether this is the case. Meanwhile, these results support the potential for the *Nbd14a,b* mutant to serve as a platform for first-pass screens of D14 variants for altered SL signaling activities.

## DISCUSSION

Several assays have been used or developed to evaluate the activity of SL receptors. These can be useful tools to investigate the contributions of specific amino acids to SL recognition or signaling, or to screen for chemicals that affect SL signaling. There are three major types of assays for SL receptors: 1) those that evaluate a SL receptor alone, 2) those that test protein-protein interactions between a SL receptor and MAX2 and/or its targets, and 3) those that report degradation of the SL receptor or its SMXL target protein(s), which are direct results of SL signaling.

### *In vitro* assays for strigolactone binding and hydrolysis by D14

A range of biochemical techniques have been used to evaluate the ability of D14 or KAI2 proteins to bind, hydrolyze, and be activated by SL. Isothermal calorimetry (ITC) and surface plasmon resonance are effective ways to measure the *in vitro* affinity (i.e. K_d_) of SL receptors for SL. These techniques have been used to study D14, several KAI2-like proteins in *Physcomitrella patens*, and a set of 60 mutants of ShHTL7 (Kagiyama et al., 2013; Bürger et al., 2019; Pang et al., 2020). Yoshimulactone green (YLG) competition assays are another popular method to assess the affinity of a SL receptor for different ligands. In these *in vitro* assays, the SL analog YLG is hydrolyzed by the receptor, releasing a fluorescent byproduct. Compounds are tested for their ability to competitively interfere with YLG hydrolysis, producing a half-maximal inhibitory concentration (IC_50_) value that generally corresponds with the receptor’s affinity for the compound (Tsuchiya et al., 2015). The YLG competition assay was used to identify a D14 inhibitor from a chemical library of 800 compounds, as well as to test a set of binding pocket mutants of ShHTL7 (Uraguchi et al., 2018; Yoshimura et al., 2018). Hydrolysis of SL results in attachment of the methylbutenolide group from SL onto the catalytic His residue of the receptor (de Saint Germain et al., 2016; Yao et al., 2016). Formation of this “covalently linked intermediate molecule” (CLIM) can be tracked with liquid chromatography-mass spectrometry, and serves as another readout of a SL receptor’s activity on a SL or SL analog. However, substrate-binding and hydrolysis rates may correlate poorly with the receptor’s signaling activity (Uraguchi et al., 2018). Differential scanning fluorimetry (DSF) or nano differential scanning fluorimetry have often been used to monitor shifts in the melting temperature or intrinsic fluorescence of SL receptors that are induced by potential ligands *in vitro* (Hamiaux et al., 2012; Waters et al., 2015; Hamiaux et al., 2018; Bürger et al., 2019; Seto et al., 2019). These changes may indicate conformational changes in the SL receptor that correspond with its activation for downstream signal transduction.

*In vitro* assays for SL-binding or activation of the SL receptor require purification of the protein of interest, which does not seem to pose a significant roadblock. A bigger issue is that these assays do not report how SL perception by the receptor affects downstream signaling events or incorporate the effects of *in vivo* factors (e.g. protein partners) on the receptor’s ligand-binding, hydrolysis, or signaling activities. For example, the presence of MAX2 is known to slow SL hydrolysis by D14 *in vitro* and the affinity of D14 for D53/SMXL7 is enhanced by the presence of MAX2 (Yao et al., 2016; Shabek et al., 2018). This is also a potential weakness of *in silico* approaches such as molecular docking, pharmacophore modeling, and molecular dynamics simulations that model interactions between SL receptors in isolation and potential ligands (Mashita et al., 2016; Fukui et al., 2017; Bürger and Chory, 2020; Lee et al., 2020).

### An *in vivo* assay for strigolactone binding

A recent study has provided an exciting new approach to measure SL binding *in vivo* as well as *in vitro*. These SL biosensors incorporate a circularly-permuted GFP (cpGFP) protein into an external loop joining alpha-helices 6 and 7 of DAD2 or ShHTL7 (Chesterfield et al., 2020). SL induces conformational shifts in DAD2 or ShHTL7 that also affect the conformation of cpGFP, reducing its fluorescence. A second fluorescent protein fused to the biosensor enables ratiometric measurements of fluorescence that bypass problems with varying biosensor abundance. In tobacco protoplasts, the DAD2-based biosensor can detect *rac*-GR24 concentrations as low as 100 nM (Chesterfield et al., 2020). Because this system is sensitive to even single amino acid shifts in the placement of the cpGFP and must be fine-tuned for each SL receptor, it may be better suited for screening for agonist or antagonist molecules of the receptor than evaluating the effects of receptor mutations. Also, while this system is able to provide valuable information on binding and detection of SLs by two SL receptors *in vitro* or *in vivo*, it does not address how perception affects downstream signalling.

### Assays for strigolactone-induced protein-protein interactions

Other SL receptor assays have focused on protein-protein interactions that are induced by SL perception. One advantage of these approaches is that they can identify factors beyond ligand-binding and hydrolysis that affect SL signaling. *In vitro* pulldowns, co-immunoprecipitation from plant tissue, size-exclusion chromatography, and AlphaScreen are proven ways to assess interactions between SL receptors, MAX2/D3, and SMXL proteins (Jiang et al., 2013; Zhou et al., 2013; Zhao et al., 2015; Yao et al., 2016; Yao et al., 2017; Shabek et al., 2018; Wang et al., 2020; Wang et al., 2021). Generally, these are low-throughput assays that can be quite challenging to perform due to difficulties in obtaining sufficient amounts of stable, soluble MAX2 and SMXL proteins. Yeast two-hybrid or three-hybrid assays that test interactions between SL receptors, MAX2, and/or SMXL proteins are better suited for testing the effects of SL receptor mutations or screening chemical libraries and have been successfully used for these purposes (Hamiaux et al., 2012; Toh et al., 2014; Nakamura et al., 2019; Seto et al., 2019; Lee et al., 2020; Wang et al., 2021). However, yeast-based interaction assays may produce false-positive results, at least for some KAI2 proteins (Yao et al., 2018; Wang et al., 2021).

### Assays for strigolactone-induced proteolysis

The method we have described in this paper falls within a third class of assays that measure SL-induced degradation of SL signaling components. These assays are relatively fast and highly specific readouts of SL signaling activity, whereas other downstream effects such as shoot branching, parasitic seed germination, or transcriptional responses may be affected by factors in addition to SLs. D14 is degraded after SL treatment in Arabidopsis and rice (Chevalier et al., 2014; Hu et al., 2017). An Arabidopsis line expressing a D14 fusion to luciferase has been developed as an *in vivo* assay for the ability of various SLs and SL analogs to induce D14 degradation (Sanchez et al., 2018). Most similar to our system is StrigoQuant, a dual-luminescent, ratiometric, SL signaling sensor (Samodelov et al., 2016). StrigoQuant expresses Renilla luciferase and a SMXL6 fusion to firefly luciferase in a single transcript. These two reporters are separated during translation due to an intervening 2A peptide. StrigoQuant has been deployed in Arabidopsis protoplasts, where it can report SMXL6 degradation induced by *rac*-GR24 or SLs at concentrations as low as 10 pM. Although achieving efficient protoplast isolation and transformation can be challenging, StrigoQuant offers the distinct advantage of being able to perform experiments with the many SL pathway mutants available for Arabidopsis. Co-expression of rice D14 with Strigoquant is able to restore SL-induced SMXL6-degradation responses to the Arabidopsis *d14* mutant, demonstrating that it is feasible to test SL receptor variants with this system (Samodelov et al., 2016).

Here we have shown that D14 proteins in *Nicotiana benthamiana* have similar functions in SL signaling and developmental control as in other angiosperms. The *Nbd14a,b* mutant shows increased axillary bud outgrowth, reduced shoot height, and altered leaf morphology (Figure 2, 3). It has also lost the ability to respond to exogenous SL analogs (Figure 4). Given the extensive studies of D14 in other plants, these results are not surprising. However, *N. benthamiana* is highly favored as a medium for analyses of transiently expressed plant proteins due to the simplicity of transformation and the robust transgene expression that can be achieved within a few days (Bally et al., 2018). Therefore, we propose that the *Nbd14a,b* mutant provides a useful genetic background to perform preliminary analyses of the SL signaling activity of D14 variants. This may enable explorations of how D14 mutations affect ligand specificity or how D14 interactions with specific SMXL protein targets are achieved.

### Limitations of assays for strigolactone-induced proteolysis in *N. benthamiana*

We propose to use this system as a complement to the techniques described above, as it comes with its own limitations. The success of this approach requires that heterologous proteins are compatible with endogenous *N. benthamiana* SL signaling components. For example, if NbMAX2 is not able to interact effectively with either the transiently co-expressed D14 or SMXL7 ratiometric reporter, the assays will fail. In cases where NbMAX2 is not compatible, it may be possible to co-express a MAX2 clone derived from the species of interest. Another constraint to this approach is the presence of endogenous SLs that may activate some transiently expressed SL receptors prior to the application of an agonist. Receptors with high sensitivity to SL may still be identified by causing low SMXL7 reporter abundance pre-treatment. However, adding a mutation that blocks SL biosynthesis to the *Nbd14a,b* line would be a useful way to eliminate background activation of D14 transgenes. Finally, it should be noted that our system is not suitable for studying the SL receptors in parasitic plants that mediate host perception. These SL receptors are neofunctionalized paralogs of KAI2 that target SMAX1 for degradation (Nelson, 2021). Thus, the endogenous KAI2 proteins that remain present in *Nbd14a,b* may confuse evaluations of SMAX1 reporter degradation by parasite SL receptors.

## METHODS

### Genes

*NbD14a* is found on Sol Genomics Network (SGN) scaffold Niben101Scf02153 (*N. benthamiana* Genome v1.0.1; (Bombarely et al., 2012)) and on *Nicotiana benthamiana* Sequencing Consortium (NbSC) scaffold NbLab330C11 (*N*.*benthamiana* v3.3; (Naim et al., 2012)). *NbD14b* is found on SGN scaffold Niben101Scf06949 and on NbSC scaffold NbLab330C03. *AtD14* (AT3G03990) and *AtSMXL7* (AT2G29970) have been previously described (Waters et al., 2012; Stanga et al., 2013).

### Construction of CRISPR-Cas9 constructs

20-nt guide sequences were selected from the CRISPR-P 2.0 database for the *Nicotiana benthamiana* genome (v0.4.4) to simultaneously target NbS00019774g0007 and NbS00024870g0006 with no mismatches (Bombarely et al., 2012; Liu et al., 2017). The next most likely off-target sites (based on off-score) for each guide selected had at least three mismatches and were located in intergenic or intron regions. The two guide sequences were cloned into the pHEE401E vector according to the simplified protocol for two gRNA expression cassettes for dicots (Wang et al., 2015b). Briefly, high-fidelity PCR amplification of a pCBC-DT1T2 template (containing a U6-26 terminator and U6-29 promoter) with overlapping primers was performed to add on a guide sequence and BsaI restriction site at each end. Primer sequences for NbD14-DT1-BsF, NbD14-DT1-F0, NbD14-DT2-R0, and NbD14-DT2-BsR are described in Supplementary Table 1. After purification of the extended PCR fragment, GoldenGate cloning with BsaI and pHEE401E was performed. Electrocompetent *E. coli* (strain DH5a) was transformed with the reaction product and selected on solid Luria Broth (LB) medium supplemented with 50 mg/L kanamycin. Colony PCR was performed with U6-26p-F and U6-29p-R. Plasmids were purified from colonies positive for a successful vector insertion (726 bp product) and both guide sequences were verified by Sanger sequencing with U6-26p-F and U6-29p-F. *Agrobacterium tumefaciens* (strain GV3101) was transformed with the pHEE401E-NbD14 construct and selected on solid LB medium supplemented with 50 mg/L kanamycin, 25 mg/L gentamicin, and 25 mg/L rifampicin.

### Stable transformation of *Nicotiana benthamiana*

Transformation of *N. benthamiana* with the pHEE401E-NbD14 construct was performed by the Plant Transformation Facility at University of California, Davis. Newly expanded leaves from *in vitro*-grown *Nicotiana benthamiana* plantlets were removed and cut into 1 cm^2^ pieces while suspended in a solution of *Agrobacterium tumefaciens* adjusted to an OD_600_ of 0.1-0.2. Leaf pieces were transferred abaxial side down onto co-cultivation medium consisting of Murashige and Skoog minimal organics with 8% (w/v) agar medium (MSO) supplemented with 30 g/L sucrose, 2.0 mg/L 6-benzylaminopurine (BAP) and 200 µM acetosyringone, pH 5.6-5.8, and incubated 2 d at 23°C in the dark. After 2 d, leaf pieces were transferred to shoot induction medium consisting of MSO medium supplemented with 30 g/L sucrose, 2.0 mg/L BAP, 400 mg/L carbenicillin, 250 mg/L cefotaxime, 25 mg/L hygromycin and incubated for 10 d at 26 C under a 16 h light (intensity 30 µM m^2^ s^-1^):8 h dark photoperiod. After 10 d, tissue was subcultured every 21 d onto the same medium formulation until buds developed. Developing buds were then transferred to elongation medium consisting of MSO medium supplemented with 30 g/L sucrose, 0.1 mg/L BAP, 400 mg/L carbenicillin, 250 mg/L cefotaxime, and 25 mg/L hygromycin and the tissue was subcultured every 21 d until shoots developed. After shoots reached 3-4 cm in height, they were harvested and transferred to rooting medium consisting of 0.5x MSO medium supplemented with 15 g/L sucrose, 0.2 mg/L indole-3-butyric acid (IBA), 400 mg/L carbenicillin, 250 mg/L cefotaxime, and 25 mg/L hygromycin for 14 d.

### Plant growth conditions

Plants used in growth, branching, and leaf morphology assays were grown on soil in a greenhouse in Riverside, CA from beginning of October 2019 through to mid December 2019. Plants were watered regularly every other day. Plants used in images for Figure 2A and SMXL7 degradation assays were grown on soil in a growth room under long day conditions (16 h white light at intensity 120 µM m^2^ s^-1^/8 h dark) at 22°C.

### Genotyping

DNA was extracted from young leaf tissue using the DNAzol protocol (Molecular Research Center, Inc) and analyzed by PCR with Taq DNA polymerase (New England Biolabs). *NbD14a* was amplified with NbD14a,b-F and NbD14a-3’UTR-R primers with the following thermal cycling conditions: 95°C for 3 min; 40 cycles of 95°C for 30 s, 59°C for 30 s, 68°C for 2 min; 68°C for 5 min. NbD14a,b-F and NbD14a-R were used for Sanger sequencing of purified PCR products. The first exon of *NbD14b* was amplified with NbD14a,b-F and NbD14b-Intron-R, and the second exon was amplified with NbD14b-Intron-F and NbD14b-R with the following thermal cycling conditions: 95°C for 3 min; 35 cycles of 95°C for 30 s, 52°C for 30 s, 68°C for 1 min; 68°C for 5 min. NbD14a,b-F, NbD14b-Intron-R, NbD14b-Intron-F, and NbD14b-R were used for Sanger sequencing of purified PCR products. The absence of the pHEE401E T-DNA was validated using pHEE401EhygB-F and pHEE401EhygB-R primers for the hygromycin resistance gene with the following thermal cycling conditions: 95°C for 3 min; 40 cycles of 95°C for 30 s, 57°C for 30 s, 68°C for 45 s; 68°C for 5 min.

### Plant growth, shoot branching, and leaf morphology assays

One-week old seedlings were transplanted and grown for 51 d in the greenhouse. From the base of each plant and in developmental order, each branch at the base of the node was cut off and measured from the cutoff point to the meristematic zone and the leaf was photographed. Each plant was measured from the soil level to the shoot apical meristem. Excised leaves were photographed and petiole length, leaf blade length, and leaf blade width were measured using ImageJ (NIH). Graphs and statistical analysis were performed in Prism (GraphPad).

### SMXL7 degradation assays

GR24^5DS^, GR24^ent-5DS^, and *rac*-GR24 were synthesized and purified by Dr. Adrian Scaffidi and Dr. Gavin Flematti (University of Western Australia). The ratiometric reporter for AtSMXL7, pRATIO3212-SMXL7, is previously described (Khosla et al., 2020a). pRATIO3212-SMXL7 was transformed into an *Agrobacterium tumefaciens* strain GV3101 that also carries a plasmid expressing p19, a suppressor of gene-silencing. AtD14 variants were synthesized (Twist Bioscience), cloned into an ampicillin-resistant pDONR220 entry vector, sequence-verified, and cloned into the plant transformation vector pGWB415 (Nakagawa et al., 2007) using Gateway BP and LR cloning enzymes (Invitrogen). pGWB415-D14 vectors were transformed into *A. tumefaciens* strain GV3101. Transient transformation of *Nicotiana benthamiana* and measurement of mScarlet-I and Venus was performed as described previously in a detailed protocol (Khosla and Nelson, 2020), with the following modifications for the cell densities of *A. tumefaciens* cultures resuspended in infiltration media prior to injection in *N. benthamiana* leaves: Figure 4A, final OD_600_=0.6; Figure 4B, final OD_600_=1.2, with a 1:1 composition of pRATIO/p19:pGWB415 strains; Figure 5, final OD_600_=0.9, with an 8:1 composition of pRATIO/p19:pGWB415 strains. Graphs and statistical analysis were performed in Prism (GraphPad).

## SUPPORTING INFORMATION

Table S1. Primers used in this study

## ACKNOWLEDGMENTS

We gratefully acknowledge funding support from the US National Science Foundation (NSF) Research Traineeship (NRT) Program Grant DGE-1922642 “Plants3D” to AW, and NSF grants IOS-1740560 and IOS-1856741 to DCN. We thank Dr. Gavin Flematti and Dr. Adrian Scaffidi (University of Western Australia) for supplying *rac*-GR24 and purified GR24 enantiomers. We thank James Eckhardt and Claudia Sepulveda for providing assistance at early stages of the project. We also thank Dr. David Tricoli and Bailey Van Bockern at the Plant Transformation Facility of the University of California, Davis for performing transformation of *N. benthamiana*.

## AUTHOR CONTRIBUTIONS

Experiments were designed, carried out, and analyzed by ARFW, JAM, AK, and DCN. Figure preparation by ARFW. Manuscript preparation by ARFW and DCN with contributions and final approval from all authors. Project design by DCN. Funding to support the project was secured by DCN.

## REFERENCES

Abe S, Sado A, Tanaka K, Kisugi T, Asami K, Ota S, Kim HI, Yoneyama K, Xie X, Ohnishi T, et al (2014) Carlactone is converted to carlactonoic acid by MAX1 in Arabidopsis and its methyl ester can directly interact with AtD14 in vitro. Proc Natl Acad Sci U S A 111: 18084–18089

Agusti J, Herold S, Schwarz M, Sanchez P, Ljung K, Dun EA, Brewer PB, Beveridge CA, Sieberer T, Sehr EM, et al (2011) Strigolactone signaling is required for auxin-dependent stimulation of secondary growth in plants. Proc Natl Acad Sci U S A 108: 20242–20247

Ahmad MZ, Rehman NU, Yu S, Zhou Y, Haq BU, Wang J, Li P, Zeng Z, Zhao J (2020) GmMAX2-D14 and -KAI interaction-mediated SL and KAR signaling play essential roles in soybean root nodulation. Plant J 101: 334–351

Akiyama K, Matsuzaki K-I, Hayashi H (2005) Plant sesquiterpenes induce hyphal branching in arbuscular mycorrhizal fungi. Nature 435: 824–827

Arite T, Umehara M, Ishikawa S, Hanada A, Maekawa M, Yamaguchi S, Kyozuka J (2009) d14, a Strigolactone-Insensitive Mutant of Rice, Shows an Accelerated Outgrowth of Tillers. Plant and Cell Physiology 50: 1416–1424

Arumingtyas EL, Floyd RS, Gregory MJ, Mufert IC (1992) Branching in Pisum: inheritance and allelism tests with 17 ramosus mutants. Pisum Genetics 24: 17–31

Bally J, Jung H, Mortimer C, Naim F, Philips JG, Hellens R, Bombarely A, Goodin MM, Waterhouse PM (2018) The Rise and Rise of Nicotiana benthamiana: A Plant for All Reasons. Annu Rev Phytopathol 56: 405–426

Besserer A, Bécard G, Jauneau A, Roux C, Séjalon-Delmas N (2008) GR24, a synthetic analog of strigolactones, stimulates the mitosis and growth of the arbuscular mycorrhizal fungus Gigaspora rosea by boosting its energy metabolism. Plant Physiol 148: 402–413

Besserer A, Puech-Pagès V, Kiefer P, Gomez-Roldan V, Jauneau A, Roy S, Portais J-C, Roux C, Bécard G, Séjalon-Delmas N (2006) Strigolactones stimulate arbuscular mycorrhizal fungi by activating mitochondria. PLoS Biol 4: e226

Bombarely A, Rosli HG, Vrebalov J, Moffett P, Mueller LA, Martin GB (2012) A draft genome sequence of Nicotiana benthamiana to enhance molecular plant-microbe biology research. Mol Plant Microbe Interact 25: 1523–1530

Bouwmeester H, Li C, Thiombiano B, Rahimi M, Dong L (2021) Adaptation of the parasitic plant lifecycle: germination is controlled by essential host signaling molecules. Plant Physiol 185: 1292–1308

Bunsick M, Toh S, Wong C, Xu Z, Ly G, McErlean CSP, Pescetto G, Nemrish KE, Sung P, Li JD, et al (2020) SMAX1-dependent seed germination bypasses GA signalling in Arabidopsis and Striga. Nat Plants 6: 646–652

Bu Q, Lv T, Shen H, Luong P, Wang J, Wang Z, Huang Z, Xiao L, Engineer C, Kim TH, et al (2014) Regulation of drought tolerance by the F-box protein MAX2 in Arabidopsis. Plant Physiol 164: 424–439

Bürger M, Chory J (2020) In-silico analysis of the strigolactone ligand-receptor system. Plant Direct 4: e00263

Bürger M, Mashiguchi K, Lee HJ, Nakano M, Takemoto K, Seto Y, Yamaguchi S, Chory J (2019) Structural Basis of Karrikin and Non-natural Strigolactone Perception in Physcomitrella patens. Cell Rep 26: 855–865.e5

Carbonnel S, Das D, Varshney K, Kolodziej MC, Villaécija-Aguilar JA, Gutjahr C (2020) The karrikin signaling regulator SMAX1 controls Lotus japonicus root and root hair development by suppressing ethylene biosynthesis. Proc Natl Acad Sci U S A 117: 21757–21765

Carbonnel S, Torabi S, Griesmann M, Bleek E, Tang Y (2019) Duplicated KAI2 receptors with divergent ligand-binding specificities control distinct developmental traits in Lotus japonicus. bioRxiv

Chesterfield RJ, Whitfield JH, Pouvreau B, Cao D, Alexandrov K, Beveridge CA, Vickers CE (2020) Rational Design of Novel Fluorescent Enzyme Biosensors for Direct Detection of Strigolactones. ACS Synth Biol 9: 2107–2118

Chevalier F, Nieminen K, Sánchez-Ferrero JC, Rodríguez ML, Chagoyen M, Hardtke CS, Cubas P (2014) Strigolactone promotes degradation of DWARF14, an α/β hydrolase essential for strigolactone signaling in Arabidopsis. Plant Cell 26: 1134–1150

Choi J, Lee T, Cho J, Servante EK, Pucker B, Summers W, Bowden S, Rahimi M, An K, An G, et al (2020) The negative regulator SMAX1 controls mycorrhizal symbiosis and strigolactone biosynthesis in rice. Nat Commun 11: 2114

Conn CE, Bythell-Douglas R, Neumann D, Yoshida S, Whittington B, Westwood JH, Shirasu K, Bond CS, Dyer KA, Nelson DC (2015) PLANT EVOLUTION. Convergent evolution of strigolactone perception enabled host detection in parasitic plants. Science 349: 540–543

Drummond RSM, Sheehan H, Simons JL, Martínez-Sánchez NM, Turner RM, Putterill J, Snowden KC (2011) The Expression of Petunia Strigolactone Pathway Genes is Altered as Part of the Endogenous Developmental Program. Front Plant Sci 2: 115

Flematti GR, Scaffidi A, Waters MT, Smith SM (2016) Stereospecificity in strigolactone biosynthesis and perception. Planta 243: 1361–1373

Fukui K, Yamagami D, Ito S, Asami T (2017) A Taylor-Made Design of Phenoxyfuranone-Type Strigolactone Mimic. Front Plant Sci 8: 936

Gomez-Roldan V, Fermas S, Brewer PB, Puech-Pagès V, Dun EA, Pillot J-P, Letisse F, Matusova R, Danoun S, Portais J-C, et al (2008) Strigolactone inhibition of shoot branching. Nature 455: 189–194

Gutjahr C, Gobbato E, Choi J, Riemann M, Johnston MG, Summers W, Carbonnel S, Mansfield C, Yang S-Y, Nadal M, et al (2015) Rice perception of symbiotic arbuscular mycorrhizal fungi requires the karrikin receptor complex. Science 350: 1521–1524

Hamiaux C, Drummond RSM, Janssen BJ, Ledger SE, Cooney JM, Newcomb RD, Snowden KC (2012) DAD2 is an α/β hydrolase likely to be involved in the perception of the plant branching hormone, strigolactone. Curr Biol 22: 2032–2036

Hamiaux C, Drummond RSM, Luo Z, Lee HW, Sharma P, Janssen BJ, Perry NB, Denny WA, Snowden KC (2018) Inhibition of strigolactone receptors by N-phenylanthranilic acid derivatives: Structural and functional insights. J Biol Chem 293: 6530–6543

Hu Q, He Y, Wang L, Liu S, Meng X, Liu G, Jing Y, Chen M, Song X, Jiang L, et al (2017) DWARF14, A Receptor Covalently Linked with the Active Form of Strigolactones, Undergoes Strigolactone-Dependent Degradation in Rice. Front Plant Sci 8: 1935

Jiang L, Liu X, Xiong G, Liu H, Chen F, Wang L, Meng X, Liu G, Yu H, Yuan Y, et al (2013) DWARF 53 acts as a repressor of strigolactone signalling in rice. Nature 504: 401–405

Kagiyama M, Hirano Y, Mori T, Kim S-Y, Kyozuka J, Seto Y, Yamaguchi S, Hakoshima T (2013) Structures of D14 and D14L in the strigolactone and karrikin signaling pathways. Genes Cells 18: 147–160

Kalliola M, Jakobson L, Davidsson P, Pennanen V, Waszczak C, Yarmolinsky D, Zamora O, Palva ET, Kariola T, Kollist H, et al (2020) Differential role of MAX2 and strigolactones in pathogen, ozone, and stomatal responses. Plant Direct 4: e00206

Kapulnik Y, Delaux P-M, Resnick N, Mayzlish-Gati E, Wininger S, Bhattacharya C, Séjalon-Delmas N, Combier J-P, Bécard G, Belausov E, et al (2011) Strigolactones affect lateral root formation and root-hair elongation in Arabidopsis. Planta 233: 209–216

Khosla A, Morffy N, Li Q, Faure L, Chang SH, Yao J, Zheng J, Cai ML, Stanga JP, Flematti GR, et al (2020a) Structure-Function Analysis of SMAX1 Reveals Domains that Mediate its Karrikin-Induced Proteolysis and Interaction with the Receptor KAI2. The Plant Cell tpc.00752.2019

Khosla A, Nelson DC (2016) Strigolactones, super hormones in the fight against Striga. Curr Opin Plant Biol 33: 57–63

Khosla A, Nelson DC (2020) Ratiometric Measurement of Protein Abundance after Transient Expression of a Transgene in Nicotiana benthamiana. Bio Protoc 10: e3747

Khosla A, Rodriguez-Furlan C, Kapoor S, Van Norman JM, Nelson DC (2020b) A series of dual-reporter vectors for ratiometric analysis of protein abundance in plants. Plant Direct. doi: 10.1002/pld3.231

Kobae Y, Kameoka H, Sugimura Y, Saito K, Ohtomo R, Fujiwara T, Kyozuka J (2018) Strigolactone Biosynthesis Genes of Rice are Required for the Punctual Entry of Arbuscular Mycorrhizal Fungi into the Roots. Plant Cell Physiol 59: 544– 553

Kretzschmar T, Kohlen W, Sasse J, Borghi L, Schlegel M, Bachelier JB, Reinhardt D, Bours R, Bouwmeester HJ, Martinoia E (2012) A petunia ABC protein controls strigolactone-dependent symbiotic signalling and branching. Nature 483: 341–344

Lahari Z, Ullah C, Kyndt T, Gershenzon J, Gheysen G (2019) Strigolactones enhance root-knot nematode (Meloidogyne graminicola) infection in rice by antagonizing the jasmonate pathway. New Phytol 224: 454–465

Lauressergues D, André O, Peng J, Wen J, Chen R, Ratet P, Tadege M, Mysore KS, Rochange SF (2015) Strigolactones contribute to shoot elongation and to the formation of leaf margin serrations in Medicago truncatula R108. J Exp Bot 66: 1237–1244

Lee HW, Sharma P, Janssen BJ, Drummond RSM, Luo Z, Hamiaux C, Collier T, Allison JR, Newcomb RD, Snowden KC (2020) Flexibility of the petunia strigolactone receptor DAD2 promotes its interaction with signaling partners. J Biol Chem 295: 4181–4193

Liu H, Ding Y, Zhou Y, Jin W, Xie K, Chen L-L (2017) CRISPR-P 2.0: An Improved CRISPR-Cas9 Tool for Genome Editing in Plants. Mol Plant 10: 530–532

Liu Q, Zhang Y, Matusova R, Charnikhova T, Amini M, Jamil M, Fernandez-Aparicio M, Huang K, Timko MP, Westwood JH, et al (2014) Striga hermonthica MAX2 restores branching but not the Very Low Fluence Response in the Arabidopsis thaliana max2 mutant. New Phytol 202: 531–541

Liu R, Hou J, Li H, Xu P, Zhang Z, Zhang X (2021) Association of TaD14-4D, a Gene Involved in Strigolactone Signaling, with Yield Contributing Traits in Wheat. Int J Mol Sci. doi: 10.3390/ijms22073748

Li W, Nguyen KH, Chu HD, Van Ha C, Watanabe Y, Osakabe Y, Leyva-González MA, Sato M, Toyooka K, Voges L, et al (2017) The karrikin receptor KAI2 promotes drought resistance in Arabidopsis thaliana. PLoS Genet 13: e1007076

Li W, Nguyen KH, Chu HD, Watanabe Y, Osakabe Y, Sato M, Toyooka K, Seo M, Tian L, Tian C, et al (2020) Comparative functional analyses of DWARF14 and KARRIKIN INSENSITIVE 2 in drought adaptation of Arabidopsis thaliana. Plant J 103: 111–127

Luke G, Escuin H, De Felipe P, Ryan M (2010) 2A to the fore - research, technology and applications. Biotechnol Genet Eng Rev 26: 223–260

Machin DC, Hamon-Josse M, Bennett T (2020) Fellowship of the rings: a saga of strigolactones and other small signals. New Phytol 225: 621–636

Marzec M, Gruszka D, Tylec P, Szarejko I (2016) Identification and functional analysis of the HvD14 gene involved in strigolactone signaling in Hordeum vulgare. Physiol Plant 158: 341–355

Mashita O, Koishihara H, Fukui K, Nakamura H, Asami T (2016) Discovery and identification of 2-methoxy-1-naphthaldehyde as a novel strigolactone-signaling inhibitor. J Pestic Sci 41: 71–78

Naim F, Nakasugi K, Crowhurst RN, Hilario E, Zwart AB, Hellens RP, Taylor JM, Waterhouse PM, Wood CC (2012) Advanced engineering of lipid metabolism in Nicotiana benthamiana using a draft genome and the V2 viral silencing-suppressor protein. PLoS One 7: e52717

Nakagawa T, Kurose T, Hino T, Tanaka K, Kawamukai M, Niwa Y, Toyooka K, Matsuoka K, Jinbo T, Kimura T (2007) Development of series of gateway binary vectors, pGWBs, for realizing efficient construction of fusion genes for plant transformation. J Biosci Bioeng 104: 34–41

Nakamura H, Hirabayashi K, Miyakawa T, Kikuzato K, Hu W, Xu Y, Jiang K, Takahashi I, Niiyama R, Dohmae N, et al (2019) Triazole Ureas Covalently Bind to Strigolactone Receptor and Antagonize Strigolactone Responses. Molecular Plant 12: 44–58

Napoli CA, Ruehle J (1996) New Mutations Affecting Meristem Growth and Potential in Petunia hybridaVilm. J Hered 87: 371–377

Nasir F, Tian L, Shi S, Chang C, Ma L, Gao Y, Tian C (2019) Strigolactones positively regulate defense against Magnaporthe oryzae in rice (Oryza sativa). Plant Physiol Biochem 142: 106–116

Nelson DC (2021) The mechanism of host-induced germination in root parasitic plants. Plant Physiol 185: 1353–1373

Nelson DC, Flematti GR, Riseborough J-A, Ghisalberti EL, Dixon KW, Smith SM (2010) Karrikins enhance light responses during germination and seedling development in Arabidopsis thaliana. Proc Natl Acad Sci U S A 107: 7095–7100

Nelson DC, Riseborough J-A, Flematti GR, Stevens J, Ghisalberti EL, Dixon KW, Smith SM (2009) Karrikins discovered in smoke trigger Arabidopsis seed germination by a mechanism requiring gibberellic acid synthesis and light. Plant Physiol 149: 863–873

Nelson DC, Scaffidi A, Dun EA, Waters MT, Flematti GR, Dixon KW, Beveridge CA, Ghisalberti EL, Smith SM (2011) F-box protein MAX2 has dual roles in karrikin and strigolactone signaling in Arabidopsis thaliana. Proc Natl Acad Sci U S A 108: 8897–8902

Pang Z, Zhang X, Ma F, Liu J, Zhang H, Wang J, Wen X, Xi Z (2020) Comparative Studies of Potential Binding Pocket Residues Reveal the Molecular Basis of ShHTL Receptors in the Perception of GR24 in Striga. J Agric Food Chem 68: 12729– 12737

Ruyter-Spira C, Kohlen W, Charnikhova T, van Zeijl A, van Bezouwen L, de Ruijter N, Cardoso C, Lopez-Raez JA, Matusova R, Bours R, et al (2011) Physiological effects of the synthetic strigolactone analog GR24 on root system architecture in Arabidopsis: another belowground role for strigolactones? Plant Physiol 155: 721– 734

de Saint Germain A, Clavé G, Badet-Denisot M-A, Pillot J-P, Cornu D, Le Caer J-P, Burger M, Pelissier F, Retailleau P, Turnbull C, et al (2016) An histidine covalent receptor and butenolide complex mediates strigolactone perception. Nat Chem Biol 12: 787–794

de Saint Germain A, Jacobs A, Brun G, Pouvreau J-B, Braem L, Cornu D, Clavé G, Baudu E, Steinmetz V, Servajean V, et al (2020) A Phelipanche ramosa KAI2 Protein Perceives enzymatically Strigolactones and Isothiocyanates. bioRxiv 2020.06.09.136473

Samodelov SL, Beyer HM, Guo X, Augustin M, Jia K-P, Baz L, Ebenhöh O, Beyer P, Weber W, Al-Babili S, et al (2016) StrigoQuant: A genetically encoded biosensor for quantifying strigolactone activity and specificity. Sci Adv 2: e1601266

Sanchez E, Artuso E, Lombardi C, Visentin I, Lace B, Saeed W, Lolli ML, Kobauri P, Ali Z, Spyrakis F, et al (2018) Structure-activity relationships of strigolactones via a novel, quantitative in planta bioassay. J Exp Bot 69: 2333–2343

Scaffidi A, Waters MT, Ghisalberti EL, Dixon KW, Flematti GR, Smith SM (2013) Carlactone-independent seedling morphogenesis in Arabidopsis. Plant J 76: 1–9

Scaffidi A, Waters MT, Sun YK, Skelton BW, Dixon KW, Ghisalberti EL, Flematti GR, Smith SM (2014) Strigolactone Hormones and Their Stereoisomers Signal through Two Related Receptor Proteins to Induce Different Physiological Responses in Arabidopsis. Plant Physiol 165: 1221–1232

Seto Y, Yasui R, Kameoka H, Tamiru M, Cao M, Terauchi R, Sakurada A, Hirano R, Kisugi T, Hanada A, et al (2019) Strigolactone perception and deactivation by a hydrolase receptor DWARF14. Nat Commun 10: 191

Shabek N, Ticchiarelli F, Mao H, Hinds TR, Leyser O, Zheng N (2018) Structural plasticity of D3-D14 ubiquitin ligase in strigolactone signalling. Nature 563: 652–656

Shen H, Luong P, Huq E (2007) The F-box protein MAX2 functions as a positive regulator of photomorphogenesis in Arabidopsis. Plant Physiol 145: 1471–1483

Shindo M, Yamamoto S, Shimomura K, Umehara M (2020) Strigolactones Decrease Leaf Angle in Response to Nutrient Deficiencies in Rice. Front Plant Sci 11: 135

Shinohara N, Taylor C, Leyser O (2013) Strigolactone can promote or inhibit shoot branching by triggering rapid depletion of the auxin efflux protein PIN1 from the plasma membrane. PLoS Biol 11: e1001474

Soundappan I, Bennett T, Morffy N, Liang Y, Stanga JP, Abbas A, Leyser O, Nelson DC (2015) SMAX1-LIKE/D53 Family Members Enable Distinct MAX2-Dependent Responses to Strigolactones and Karrikins in Arabidopsis. Plant Cell 27: 3143–3159

Stanga JP, Morffy N, Nelson DC (2016) Functional redundancy in the control of seedling growth by the karrikin signaling pathway. Planta 243: 1397–1406

Stanga JP, Smith SM, Briggs WR, Nelson DC (2013) SUPPRESSOR OF MORE AXILLARY GROWTH2 1 controls seed germination and seedling development in Arabidopsis. Plant Physiol 163: 318–330

Stanic M, Hickerson NMN, Arunraj R, Samuel MA (2021) Gene-editing of the strigolactone receptor BnD14 confers promising shoot architectural changes in Brassica napus (canola). Plant Biotechnol J 19: 639–641

Sun X-D, Ni M (2011) HYPOSENSITIVE TO LIGHT, an alpha/beta fold protein, acts downstream of ELONGATED HYPOCOTYL 5 to regulate seedling de-etiolation. Mol Plant 4: 116–126

Swarbreck SM, Guerringue Y, Matthus E, Jamieson FJC, Davies JM (2019) Impairment in karrikin but not strigolactone sensing enhances root skewing in Arabidopsis thaliana. Plant J 98: 607–621

Toh S, Holbrook-Smith D, Stogios PJ, Onopriyenko O, Lumba S, Tsuchiya Y, Savchenko A, McCourt P (2015) Structure-function analysis identifies highly sensitive strigolactone receptors in Striga. Science 350: 203–207

Toh S, Holbrook-Smith D, Stokes ME, Tsuchiya Y, McCourt P (2014) Detection of parasitic plant suicide germination compounds using a high-throughput Arabidopsis HTL/KAI2 strigolactone perception system. Chem Biol 21: 988–998

Tsuchiya Y, Yoshimura M, Sato Y, Kuwata K, Toh S, Holbrook-Smith D, Zhang H, McCourt P, Itami K, Kinoshita T, et al (2015) PARASITIC PLANTS. Probing strigolactone receptors in Striga hermonthica with fluorescence. Science 349: 864– 868

Ueda H, Kusaba M (2015) Strigolactone Regulates Leaf Senescence in Concert with Ethylene in Arabidopsis. Plant Physiol 169: 138–147

Umehara M, Hanada A, Yoshida S, Akiyama K, Arite T, Takeda-Kamiya N, Magome H, Kamiya Y, Shirasu K, Yoneyama K, et al (2008) Inhibition of shoot branching by new terpenoid plant hormones. Nature 455: 195–200

Uraguchi D, Kuwata K, Hijikata Y, Yamaguchi R, Imaizumi H, Am S, Rakers C, Mori N, Akiyama K, Irle S, et al (2018) A femtomolar-range suicide germination stimulant for the parasitic plant Striga hermonthica. Science 362: 1301–1305

Van Ha C, Leyva-González MA, Osakabe Y, Tran UT, Nishiyama R, Watanabe Y, Tanaka M, Seki M, Yamaguchi S, Van Dong N, et al (2014) Positive regulatory role of strigolactone in plant responses to drought and salt stress. Proc Natl Acad Sci U S A 111: 851–856

Villaécija-Aguilar JA, Hamon-Josse M, Carbonnel S, Kretschmar A, Schmid C, Dawid C, Bennett T, Gutjahr C (2019) SMAX1/SMXL2 regulate root and root hair development downstream of KAI2-mediated signalling in Arabidopsis. PLOS Genetics 15: e1008327

Wang L, Wang B, Jiang L, Liu X, Li X, Lu Z, Meng X, Wang Y, Smith SM, Li J (2015a) Strigolactone Signaling in Arabidopsis Regulates Shoot Development by Targeting D53-Like SMXL Repressor Proteins for Ubiquitination and Degradation. Plant Cell 27: 3128–3142

Wang L, Xu Q, Yu H, Ma H, Li X, Yang J, Chu J, Xie Q, Wang Y, Smith SM, et al (2020) Strigolactone and Karrikin Signaling Pathways Elicit Ubiquitination and Proteolysis of SMXL2 to Regulate Hypocotyl Elongation in Arabidopsis. Plant Cell 32: 2251–2270

Wang P, Zhang S, Qiao J, Sun Q, Shi Q, Cai C, Mo J, Chu Z, Yuan Y, Du X, et al (2019) Functional analysis of the GbDWARF14 gene associated with branching development in cotton. PeerJ 7: e6901

Wang Y, Yao R, Du X, Guo L, Chen L, Xie D, Smith SM (2021) Molecular basis for high ligand sensitivity and selectivity of strigolactone receptors in Striga. Plant Physiol 185: 1411–1428

Wang Z-P, Xing H-L, Dong L, Zhang H-Y, Han C-Y, Wang X-C, Chen Q-J (2015b) Egg cell-specific promoter-controlled CRISPR/Cas9 efficiently generates homozygous mutants for multiple target genes in Arabidopsis in a single generation. Genome Biol 16: 144

Waters MT, Gutjahr C, Bennett T, Nelson DC (2017) Strigolactone Signaling and Evolution. Annu Rev Plant Biol 68: 291–322

Waters MT, Nelson DC, Scaffidi A, Flematti GR, Sun YK, Dixon KW, Smith SM (2012) Specialisation within the DWARF14 protein family confers distinct responses to karrikins and strigolactones in Arabidopsis. Development 139: 1285–1295

Waters MT, Scaffidi A, Moulin SLY, Sun YK, Flematti GR, Smith SM (2015) A Selaginella moellendorffii Ortholog of KARRIKIN INSENSITIVE2 Functions in Arabidopsis Development but Cannot Mediate Responses to Karrikins or Strigolactones. Plant Cell 27: 1925–1944

Wen C, Xi L, Gao B, Wang K, Lv S, Kou Y, Ma N, Zhao L (2015) Roles of DgD14 in regulation of shoot branching in chrysanthemum (Dendranthema grandiflorum “Jinba”). Plant Physiol Biochem 96: 241–253

Yamada Y, Furusawa S, Nagasaka S, Shimomura K, Yamaguchi S, Umehara M (2014) Strigolactone signaling regulates rice leaf senescence in response to a phosphate deficiency. Planta 240: 399–408

Yao J, Mashiguchi K, Scaffidi A, Akatsu T, Melville KT, Morita R, Morimoto Y, Smith SM, Seto Y, Flematti GR, et al (2018) An allelic series at the KARRIKIN INSENSITIVE 2 locus of Arabidopsis thaliana decouples ligand hydrolysis and receptor degradation from downstream signalling. Plant J 96: 75–89

Yao R, Ming Z, Yan L, Li S, Wang F, Ma S, Yu C, Yang M, Chen L, Chen L, et al (2016) DWARF14 is a non-canonical hormone receptor for strigolactone. Nature 536: 469–473

Yao R, Wang F, Ming Z, Du X, Chen L, Wang Y, Zhang W, Deng H, Xie D (2017) ShHTL7 is a non-canonical receptor for strigolactones in root parasitic weeds. Cell Res 27: 838–841

Yoshimura M, Sato A, Kuwata K, Inukai Y, Kinoshita T, Itami K, Tsuchiya Y, Hagihara S (2018) Discovery of Shoot Branching Regulator Targeting Strigolactone Receptor DWARF14. ACS Cent Sci 4: 230–234

Zhao L-H, Zhou XE, Yi W, Wu Z, Liu Y, Kang Y, Hou L, de Waal PW, Li S, Jiang Y, et al (2015) Destabilization of strigolactone receptor DWARF14 by binding of ligand and E3-ligase signaling effector DWARF3. Cell Res 25: 1219–1236

Zheng J, Hong K, Zeng L, Wang L, Kang S, Qu M, Dai J, Zou L, Zhu L, Tang Z, et al (2020) Karrikin Signaling Acts Parallel to and Additively with Strigolactone Signaling to Regulate Rice Mesocotyl Elongation in Darkness. Plant Cell 32: 2780–2805

Zheng K, Wang X, Weighill DA, Guo H-B, Xie M, Yang Y, Yang J, Wang S, Jacobson DA, Guo H, et al (2016) Characterization of DWARF14 Genes in Populus. Sci Rep 6: 21593

Zhou F, Lin Q, Zhu L, Ren Y, Zhou K, Shabek N, Wu F, Mao H, Dong W, Gan L, et al (2013) D14-SCF(D3)-dependent degradation of D53 regulates strigolactone signalling. Nature 504: 406–410

