## Supplemental Table 1 for "Rapid analysis of strigolactone receptor activity in a *Nicotiana benthamiana dwarf14* mutant"

### Supplementary Table S1. Primers used in this study

| Primer name | Sequence | Use |
|---|---|---|
| NbD14-DT1-BsF | 5′-ATATATGGTCTCGATT**G**TTTGGAGAAGAGTTCAGGGGTT-3′ | gRNA cloning |
| NbD14-DT1-F0 | 5′-T**G**TTTGGAGAAGAGTTCAGGGGTTTTAGAGCTAGAAATAGC-3′ | |
| NbD14-DT2-R0 | 5'-AACTGCCATATATTTCAGACAG**C**AATCTCTTAGTCGACTCTAC-3′ | |
| NbD14-DT2-BsR | 5'-ATTATTGGTCTCGAAACTGCCATATATTTCAGACAG**C**AA-3′ | |
| U6-26p-F | 5'-TGTCCCAGGATTAGAATGATTAGGC-3' | colony PCR & sequencing of CRISPR-Cas9 construct |
| U6-29p-R | 5'-AGCCCTCTTCTTTCGATCCATCAAC-3' | |
| U6-29p-F | 5'-TTAATCCAAACTACTGCAGCCTGAC-3' | |
| pHEE401EhygB-F | 5'-CGTGCTTTCAGCTTCGATGTAG-3' | transgene detection |
| pHEE401EhygB-R | 5'-CAGTCAATGACCGCTGTTATGC-3' | |
| NbD14a,b-F | 5′-CTGAACGTACGAGTCGTAGGTTC-3′ | genotyping |
| NbD14a-3'UTR-R | 5′-TGGGAGGATATGATACGAGAGC-3′ | |
| NbD14a-R | 5′-AAGAGCTCTTCTAAGCTCTTGAGTC-3′ | |
| NbD14b-Intron-R | 5′-CGAACATCACACATCACACG-3′ | |
| NbD14b-Intron-F | 5′-TGCATCTCATTAACGGCAAG-3′ | |
| NbD14b-R | 5′-AAGAGCTCTTCTAAGCTCTTGAGCT-3′ | |
